# Large-scale structure prediction of DUF-containing protein-protein interactions

**DOI:** 10.64898/2026.08.19.745780

**Authors:** Lino Riepenhausen, Francesco Costa, Antonina Andreeva, Alex Bateman

## Abstract

**Motivation:** Continuing advances in genome and metagenome sequencing expand the number of identified conserved protein families that remain functionally uncharacterized and contain domains of unknown function (DUFs). Functional-association resources such as STRING provide biological context, but mostly do not distinguish indirect association from physical interaction. We assessed whether AlphaFold 3 complex prediction, combined with STRING evidence and domain-level analysis of interfaces and interaction partners, can help identify and characterize DUF-containing proteins.

**Results:** We generated four structural-prediction cohorts from STRING associations involving DUF-containing proteins and evaluated the predicted complexes using interface ipSAE, average pLDDT and buried surface area. An L2-regularized logistic regression model was trained on an initial cohort of predictions from high-confidence STRING associations to prioritize DUF-containing candidates likely to produce structurally confident AlphaFold 3 complexes. The model was then applied across all 12,535 organisms represented in STRING v12.0, followed by grouping into DUF-family and partner-architecture modules, covering 2,076 unique DUF families. The final L2-model screen contained 12,298 successfully modelled protein pairs, including 1,208 (9.82%) complexes meeting a strict-confidence criterion and 2,433 (19.78%) meeting a more liberal confidence criterion. Two examples suggest roles for DUF4130 in nucleic-acid-associated radical-SAM biology and DUF5819 in a bacterial system related to vitamin-K-dependent carboxylation.

**Availability and implementation:** Predicted structures and associated metadata are available through Zenodo at https://doi.org/10.5281/zenodo.21875362. The model implementation and code used to generate the analyses and figures are available at https://github.com/linoriep/Proteome-scale-structure-prediction-of-DUF-containing-protein-protein-interactions.

## 1 Introduction

The continued expansion of genome and metagenome sequencing has led to the identification of a growing number of evolutionarily conserved proteins that are uncharacterized in their function. Many of these protein families contain domains of unknown function (DUFs). DUFs are Pfam families for which a conserved domain can be recognized, but no sufficiently specific biological function has been assigned (Paysan-Lafosse *et al*. 2025). In the Pfam database, protein domains are organized into homologous sequence families, each represented by a profile hidden Markov model constructed from a curated seed alignment. Family membership reflects a common evolutionary origin and generally corresponds to a similar structural fold. 5,951 of the 25,545 families in Pfam 38.0, were classified as DUFs, being present in all organisms and most numerous in bacteria (Paysan-Lafosse *et al*. 2025). DUF-containing proteins may perform complex and biologically important functions, with genetic analyses suggesting they can be essential for cellular processes (Goodacre, Gerloff, and Uetz 2013). Because sequence conservation alone is often insufficient to determine the molecular function of DUFs, additional approaches are needed to infer their functional roles. Protein function is often relational, depending not only on protein structure, but also on interactions with other proteins and the cellular consequences of the interactions (Jacq 2001).

The STRING (Search Tool for the Retrieval of Interacting Genes/Proteins) database tries to provide this additional relational context by integrating multiple evidence channels like experimental observations, database analysis, text mining, gene neighborhood, gene fusion, phyletic co-occurrence and coexpression across organisms (Szklarczyk *et al*. 2023). However, STRING identifies possible functional associations rather than direct physical contact and incorporating structural context can strengthen these associations (Bryant, Pozzati, and Elofsson 2022, Burke *et al*. 2023).

Recent progress in deep learning for structure prediction, most notably AlphaFold, made it possible to predict structures of proteins just from their sequence (Jumper *et al*. 2021, Abramson *et al*. 2024). Further developments like AlphaFold-Multimer also enabled large-scale computational structural interaction screens and structure-informed network analysis (Humphreys *et al*. 2021, Bryant, Pozzati, and Elofsson 2022, Evans *et al*. 2022, Burke *et al*. 2023). More recently, the AlphaFold Protein Structure Database (AFDB), has also expanded beyond monomeric structures, to include protein homodimer and heterodimer complexes (Varadi *et al*. 2022, 2024, Han *et al*. 2026). However, the structural coverage of the heterodimeric structures is limited to 46 selected reference and health-priority proteomes and therefore captures only a small set of DUFs in this data. In addition, a plausibly predicted complex is not equivalent to an interaction observed *in vivo*. To further evaluate the likelihood of a genuine interaction, further metrics and biological context need to be considered (Szklarczyk *et al*. 2023).

Here we combine the functional association evidence of STRING, with AlphaFold 3 complex prediction and domain analysis, to identify structurally supported interactions involving DUFs. We first use conserved interactions to predict a set of high-confidence DUF-containing complexes and subsequently train an L2-logistic regression model, to identify plausible structurally stable interactions based on STRING data and run a large-scale AlphaFold 3 prediction screen across all 12,535 organisms represented in STRING v12.0. The resulting screen provides structurally supported functional hypotheses across many DUF families. Its utility is exemplified through the characterisation of DUF4130 and DUF5819. Our analysis shows that DUF4130 is likely involved in nucleic acid binding and S-adenosyl methionine (SAM)-dependent modification, while DUF5819 appears to have a role as part of a bacterial homolog of a Vitamin K-dependent carboxylase complex.

## 2 Methods

### STRING and Pfam

A local installation of the STRING v12.0 full scored-link database was used for the protein-protein association data (Szklarczyk *et al*. 2023). STRING combined scores range from 0 to 1000 and represent increasing confidence in a functional association. There are four combined score confidence categories, with scores ≥400 corresponding to medium confidence, scores ≥700 to high confidence and scores ≥900 to highest confidence. STRING protein identifiers were mapped to UniProt accessions using the corresponding STRING alias files and Pfam 38.0 family assignments were mapped to those accessions, to identify the domains and their residue boundaries. Protein pairs for which at least one partner could not be assigned the required protein accession were excluded from further analysis. For each pair, the two possible orientations were collapsed to one pair, as the interactions are undirected. It was possible for one of the proteins to not contain any Pfam domain architecture. A Pfam family was classified as a DUF if its short name or description contained a term beginning with “DUF” followed directly by one or more numbers (DUF[0-9]+). To reduce redundancy among related interactions, STRING interactions were grouped into modules. Each module contained interactions involving the same DUF family in one protein and the same Pfam architecture in its interaction partner. Partners without a Pfam assignment were assigned a NO_PFAM architecture. Taxonomic recurrence was defined as the number of unique NCBI/STRING taxon identifiers represented in each module. Protein functions were assigned using STRING product annotations and aliases, UniProt and reviewed Swiss-Prot protein names and function descriptions, Gene Ontology terms and Pfam domain annotations (The UniProt Consortium 2025). When these sources were unavailable, locus-tag and product-name information was used as fallback. Partner proteins were then assigned to liberal functional categories based on keyword rules applied to these annotations (**Table S3)**.

### AlphaFold 3 Predictions and Analysis

All predictions were run using a local installation of AlphaFold 3, using 1 diffusion sample per prediction, 10 recycles and seed=1. Each input contained the DUF protein as chain A and its partner as chain B. Other settings followed the official AlphaFold 3 installation (Abramson *et al*. 2024). Completed MSA (multiple sequence alignment) outputs and previously completed structure predictions were reused when available, if the same interaction was present in multiple runs. The final predicted structures were analysed with AlphaJudge v1.0.2 to obtain additional metrics including ipSAE, interface-pLDDT and pDockQ2 (Zhu *et al*. 2023, Dunbrack 2025, KosinskiLab 2026). A structure was classified as liberal confidence when ipSAE was ≥ 0.2 and average pLDDT was ≥ 70 and as strict confidence when ipSAE was ≥ 0.6 and average pLDDT was ≥ 70 (Han *et al*. 2026). Strict confidence structures were consequently a subset of liberal confidence structures. AlphaJudge scoring was only possible when there was an interface present and predictions where the scoring failed were therefore classified as low confidence. Technical failures that resulted in no AlphaFold model, such as MSA and GPU-memory failures, were excluded from the confidence-fraction calculations and not analysed further, but were retained in the reported candidate datasets. Although no explicit sequence-length cutoff was used during candidate selection, pairs over 4,600 total residues were not modelled with AlphaFold 3. The largest successfully modelled complex contained 4,587 amino-acid residues in total. To differentiate interactions where the DUF-domain was part of the interface, buried surface area (BSA) was used. The solvent-accessible surface area was calculated via the Shrake-Rupley algorithm in Biopython and BSA was derived by difference in solvent-accessible surface area between chains (Cock *et al*. 2009). BSA was calculated for all Pfam families present in each predicted complex. A DUF-domain was classified as part of the interface if the total interface area was ≥ 300 Å² and the DUF contributed ≥15% of the total buried surface area.

### Cohorts

Four cohorts of DUF-containing protein pairs were selected for AlphaFold 3 structure prediction. The first cohort used STRING’s predefined “highest confidence” cutoff and included bacterial interactions with a combined STRING score ≥900 whose corresponding module contained pairs from at least 10 bacterial taxa, resulting in 4,523 modules. The second cohort tested whether greater taxonomic recurrence could support interactions at STRING’s lower “high confidence” cutoff of a combined score ≥700. It included bacterial interactions whose corresponding module contained pairs from at least 50 taxa, resulting in 4,017 modules. For both cohorts, one pair per module was chosen based on the highest combined STRING score. In case of ties, the maximum of neighborhood, fusion, experimental and database scores was used. Preliminary analysis of the AFDB heterodimer dataset suggested that fusion evidence was associated with a higher fraction of structurally confident complexes. The third cohort therefore tested bacterial STRING interactions with a fusion channel score >1, which were grouped by the complete Pfam architectures of the two interacting proteins. Modules present in at least five organisms were retained, resulting in 2,135 modules. For each module, the pair with the highest fusion score was selected, followed by the combined STRING score in case of ties. Modules could occur in more than one cohort and were retained in each relevant cohort, but the same protein pair was predicted only once and its structure was reused for overlaps.

For a final fourth cohort, an L2-regularized logistic regression model was implemented to prioritize DUF-containing protein pairs likely to produce structurally confident predictions. It was trained on the 4,523 bacterial modules from the first cohort, with liberal confidence predictions as the positive label, an inverse regularization strength value (C) of 0.003 and class weights of 1 and 12 for negative and positive labels respectively. Different values of C and positive-class weights were evaluated by five-fold stratified cross-validation, with the combination of C=0.003 and a positive-class weight of 12 achieving the highest average precision (AP) while providing sufficient candidate coverage for the final structural screen. Missing values were median-imputed and the features were standardized. The model used the combined STRING score and all 13 individual STRING evidence channels, including direct and transferred evidence where available. All model features and fitted coefficients are provided in **Table S1**. Model performance was evaluated using five-fold stratified cross-validation. In each fold, the L2 model was fitted on the remaining four folds and evaluated on the held-out fold. Rankings based on the combined STRING score and individual STRING evidence channels were evaluated on the same held-out interactions. The model was then applied to all DUF-containing STRING pairs with a STRING combined score of ≥ 900. Pairs with a model score ≥0.5 were grouped by DUF family and partner Pfam architecture and one representative pair was selected from each group for prediction, based on the highest combined STRING score, followed by the individual evidence channels as tie-breakers. This resulted in a total of 12,374 modules for the L2-model cohort, which were predicted with AlphaFold 3. Bacteria contributed 10,422 candidate pairs, Eukaryotes 1,680 pairs and Archaea 272 pairs. **Figure 1** summarizes the selection of the four prediction cohorts and the definition of a module.

**Figure 1.**
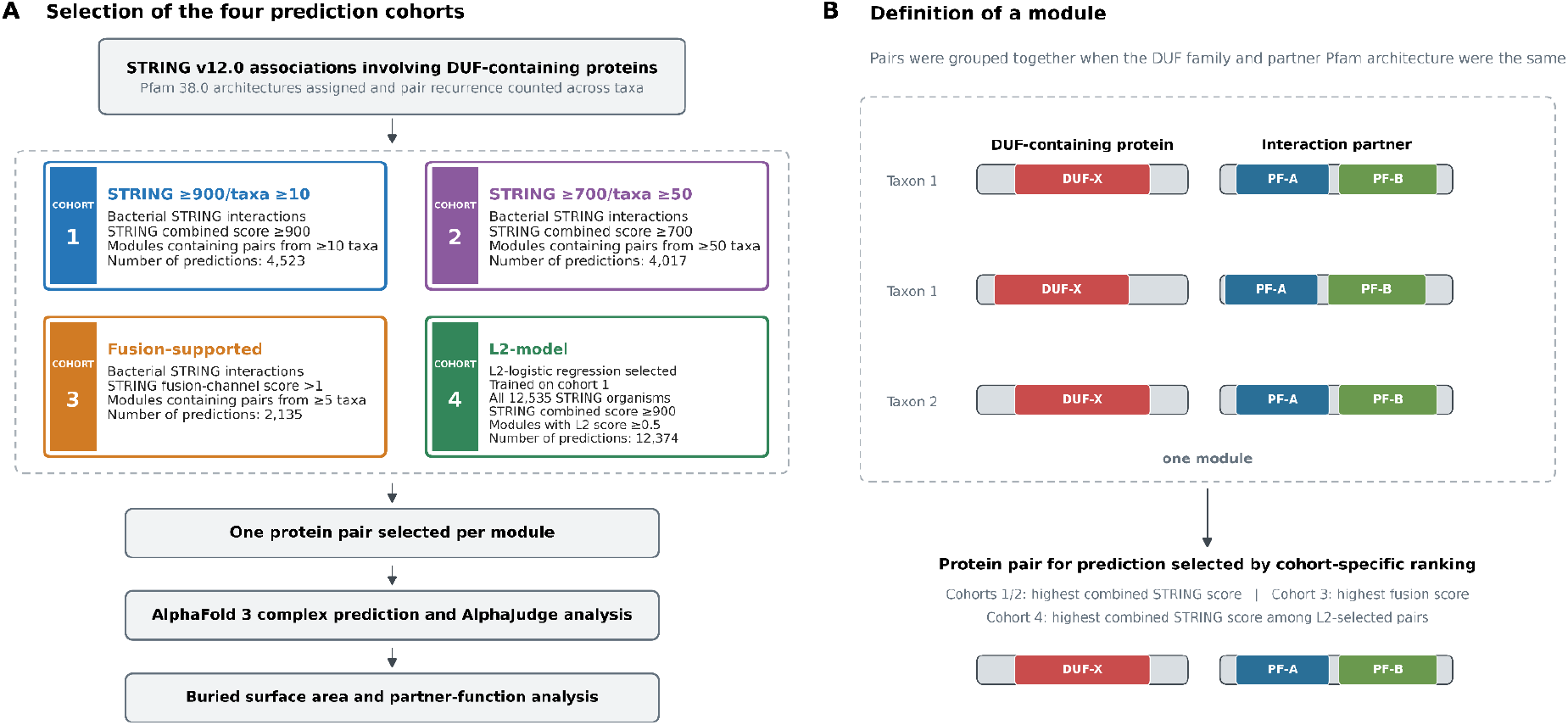
(A) Selection of the four prediction cohorts and (B) definition of a module based on DUF family and partner Pfam architecture. One protein pair was selected from each module for AlphaFold 3 prediction using cohort-specific ranking criteria. Overlapping protein pairs were predicted once and reused across cohorts. Partners without a Pfam assignment were assigned a NO_PFAM architecture.

## 3 Results

Figure 2. shows the results from all the prediction cohorts and **Table 1** shows further information about them. Structure prediction succeeded for 4,520 of 4,523 cohort one (STRING ≥900/taxa ≥10) predictions, all 4,017 cohort two (STRING ≥700/taxa ≥50) predictions, all 2,135 cohort three (fusion-supported predictions) and 12,298 of 12,374 cohort four (L2-model) predictions. The remaining predictions, which exceeded 4,600 total residues or failed because of memory limitations, were excluded from structural-confidence calculations. Across the four cohorts, 3,912 of 18,136 unique protein pairs (21.57%) occurred in more than one cohort. In the STRING ≥900/taxa ≥10 training set, the L2 model achieved an average precision (AP) of 0.372 and an area under the receiver-operating-characteristic curve (AUROC) of 0.800, compared with 0.316 and 0.781 for co-occurrence alone, 0.244 and 0.737 for neighborhood alone, and 0.142 and 0.524 for the combined STRING score alone (**Table S2**). Structural-confidence yield across the individual STRING evidence channels is shown in **Figure S1**.

**Table 1.** Candidate numbers, structural-confidence yields and DUF-family coverage across prediction cohorts and the AFDB heterodimer dataset.

| Cohort | Organisms | Successful Predictions | Liberal Confidence Fraction (%) | Strict Confidence Fraction (%) | Unique DUF Families | Unique DUF Families with $\geq 1$ Liberal-confidence member | Unique DUF Families with $\geq 1$ Strict-confidence member |
| --- | --- | --- | --- | --- | --- | --- | --- |
| AFDB heterodimer all | 46 proteomes * | 7,594,897 | 3.04 | 0.83 | 1,218 | 321 | 80 |
| AFDB heterodimer DUF-containing | 46 proteomes * | 212,084 | 1.41 | 0.37 | 1,218 | 321 | 80 |
| STRING $\geq 900$/taxa $\geq 10$ | Bacteria STRING | 4,520 | 13.58 | 7.19 | 969 | 405 | 245 |
| STRING $\geq 700$/taxa $\geq 50$ | Bacteria STRING | 4,017 | 9.58 | 4.83 | 832 | 292 | 163 |
| Fusion-supported | Bacteria STRING | 2,135 | 5.62 | 2.76 | 416 | 76 | 46 |
| L2-selected, model score $\geq 0.5$, STRING combined score $\geq 900$ | All STRING | 12,298 | 19.78 | 9.82 | 2,076 | 1,027 | 681 |
| L2-selected, model score $\geq 0.75$, STRING combined score $\geq 900$ | All STRING | 4,323 | 27.90 | 15.22 | 1,215 | 663 | 455 |
| L2-selected, model score $\geq 0.5$, STRING combined score $\geq 900$, training pairs excluded | All STRING | 9,586 | 19.11 | 9.34 | 1,944 | 859 | 548 |
| L2-selected, model score $\geq 0.75$, STRING combined score $\geq 900$, training pairs excluded | All STRING | 2,628 | 27.74 | 15.22 | 953 | 474 | 316 |

\*16 model organisms, 30 WHO global health organisms (Han *et al.* 2026)

**Figure 2.**
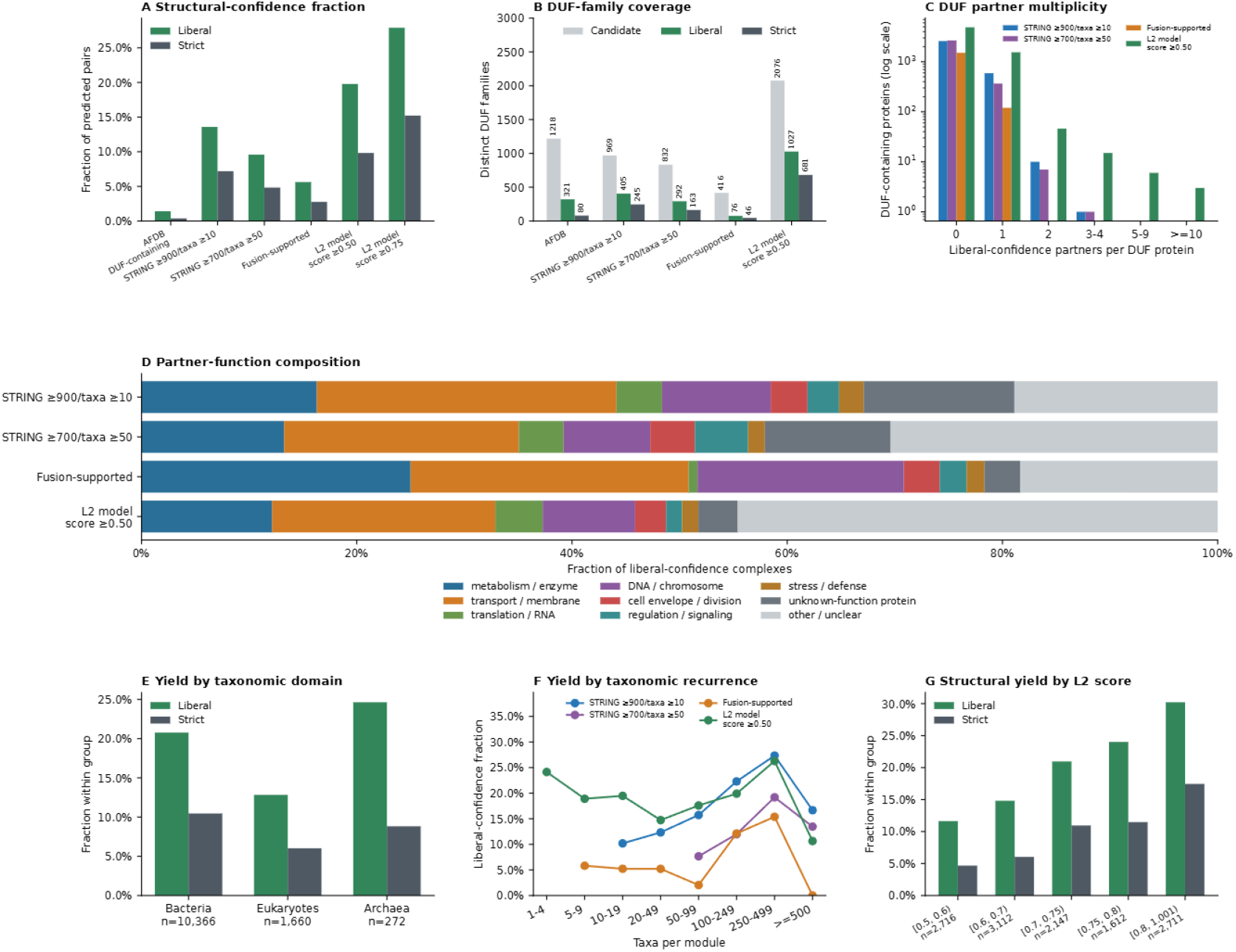
(A) Fractions of successfully predicted complexes meeting the liberal- and strict-confidence criteria, with DUF-containing AFDB heterodimers shown as a reference. (B) Numbers of distinct DUF families represented among all candidates and among complexes meeting the liberal- or strict-confidence criteria. (C) Distribution of DUF-containing proteins by the number of distinct liberal-confidence partners. (D) Functional classifications of partners in liberal-confidence complexes. (E) Liberal- and strict-confidence fractions in the bacterial,eukaryotic and archaeal subsets of the L2-selected cohort. (F) Liberal-confidence fraction according to the number of taxa in which each DUF-family/partner-architecture module was observed. (G) Liberal- and strict-confidence fractions across L2-model score intervals.

The final L2-model all-organism screen produced 2,433 liberal-confidence complexes (19.78%) and 1,208 strict-confidence complexes (9.82%). Using a stricter model-score cutoff of 0.75 retained 4,323 structures and increased the liberal-confidence fraction to 27.90% and the strict-confidence fraction to 15.22%. The bacterial STRING ≥900/taxa ≥10 cohort contained 614 liberal-confidence complexes (13.58%) and 325 strict-confidence complexes (7.19%). The STRING ≥700/taxa ≥50 cohort showed lower success, with 385 liberal confidence complexes (9.58%) and 194 strict confidence complexes (4.83%). The fusion-supported cohort had the lowest structural-confidence success, with 120 liberal confidence complexes (5.62%) and 59 strict-confidence complexes (2.76%). Among the AFDB heterodimer reference set containing at least one DUF protein, 2,992 of 212,084 complexes reached liberal confidence (1.41%) and 791 reached strict confidence (0.37%). Taxonomic recurrence showed only a weak relationship with structural confidence in the final screen.

The fraction of confident complexes in which the DUF directly contributed to the interface differed between the cohorts. DUF-interface involvement was highest in the recurrent bacterial STRING cohorts, with 421 of 614 liberal confidence structures (68.57%) in the STRING ≥900/taxa ≥10 cohort and 276 of 385 (71.69%) in the STRING ≥700/taxa ≥50 cohort passing the DUF-interface criterion. The corresponding strict-confidence fractions were 241 of 325 (74.15%) and 141 of 194 (72.68%). The fusion-supported cohort showed lower DUF-interface involvement, with 49 of 120 liberal-confidence structures (40.83%) and 22 of 59 strict confidence structures (37.29%). In the final L2-model cohort, 1,391 of 2,433 liberal confidence structures (57.17%) and 796 of 1,208 strict confidence structures (65.89%) directly involved the DUF at the interface, by the BSA criterion. Among the 1,391 liberal-confidence L2-model complexes in which the DUF formed part of the predicted interface, 649 (46.7%) contained a partner with at least one non-DUF Pfam domain and an annotation describing a specific biological or biochemical function. The corresponding subset contained 348 of 796 strict confidence DUF-interacting complexes (43.7%).

The L2 screen contained liberal-confidence members for 1,027 unique DUF families and strict-confidence members for 681 families, compared with 321 and 80 families, respectively, in the AFDB heterodimer dataset. We retrospectively matched Pfam 38.0 DUF families in the final L2-model screen to Pfam 38.2 by accession and examined predicted partner context for families that no longer met the DUF definition. Context was classified as specific when it was consistent with the later function or named biological system, as broader process when it indicated a related process without the specific function and as showing no clear relationship when relevant annotations were unavailable or absent. Of 38 identified families that were no longer classified as DUF, 15 showed specific contextual agreement, 15 showed broader agreement and 8 showed no clear relationship. Examples included XusB (formerly DUF4374) with TonB-dependent iron receptors, a levan-binding surface glycan-binding protein (formerly DUF4960) with levan-utilization proteins and the GH116 catalytic region (formerly DUF608) with beta-glucosidase partners (**Table S4**).

Genomic proximity of the predicted protein pairs in relation to structural-confidence yield was also analysed (**Figure 3**). Across all prediction cohorts, confident structures were more frequent for proteins encoded close to each other on the same locus, with the strongest effect for adjacent genes. For instance, in the STRING ≥900/taxa ≥10 cohort, 289 of 928 adjacent pairs were liberal confidence pairs (31.14%), whereas only 50 of 1,602 pairs separated by more than 1,000 locus-index units were liberal confidence pairs (3.12%). The same trend was observed in theSTRING ≥700/taxa ≥50, fusion-supported and final L2-selected cohorts, with differing strengths of the effect. It is interesting to note that the L2 selected cohort showed the least degradation in performance for distant genomic proximity.

**Figure 3.**
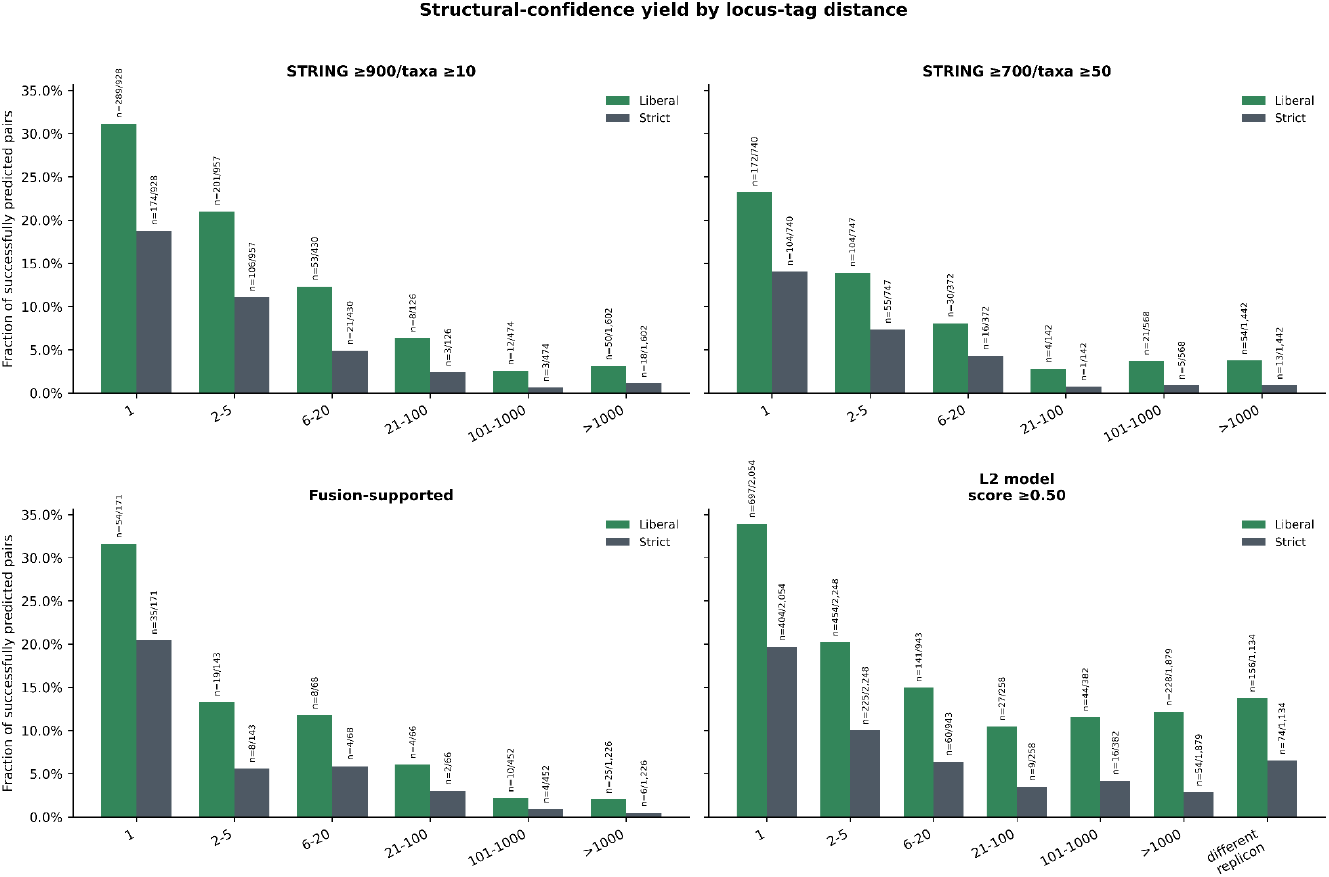
Liberal- and strict-confidence fractions are shown according to the number of genes separating each protein pair for all cohorts. Locus-tag numbering was used as a proxy for the number of separating genes.

Several predicted interactions contained sufficient structural, annotation and genomic context for further inspection, with two examples described here (**Figure 4)**. The first example was the interaction between the DUF4130 protein C806_00603 (UniProtKB: R9IYY8) and C806_00604 (UniProtKB: R9J760) from *Lachnospiraceae bacterium 3-1*. The two protein genes are adjacent and the association had a STRING combined score of 943. The predicted complex reached strict confidence, with an interface ipSAE of 0.804 and an average pLDDT of 86.3. The total interface area was 3,666 Å^2^, of which 3,252 Å^2^ was assigned to the mapped DUF4130 domain. The DUF4130 protein contains a single DUF4130 domain at residues 92-259 and its interaction partner contains a Fer4_12 domain at residues 55-179 and an HHH_3 domain at residues 327-376. Although the partner has no reviewed protein annotations, the unreviewed TrEMBL protein name (now found in UniParc) is “DNA modification/repair radical SAM protein”. The predicted structure contains a partial, distorted TIM-barrel fold, similar to those of radical SAM-enzymes and a Cys70-X3-Cys74-X2-Cys77 motif required for coordination of a [4Fe-4S] cluster (Broderick, Broderick, and Hoffman 2023). In an additional AlphaFold 3 prediction run containing SAM, the ligand was placed in the corresponding pocket next to this cysteine motif. Combination of radical SAM domain and helix-hairpin-helix (HhH_2_) domain is observed in radical S-adenosyl-l-methionine enzymes such as RlmN and Cfr involved in methylation of 23S rRNA in which the nucleic acid recognition HhH_2_ domain is located at the N-termini. Closely related to these enzymes, MiaB uses another recognition domain TRAM (OB fold) for substrate recognition. DUF4130 domains show some structural similarity (DALI Z-scores ranging from 8.8 to 6.0 (Holm and Sander 1995, Holm 2022)) to the domain arrangement DHH (PF01368)-linker domain-DHHA1 (PF02272) found in several functionally diverse proteins such as RecJ, PAP phosphatase, GAN, Cdc45 etc. This similarity is confined to the site involved in nucleotide binding, contributed by the three domains, while the similarity to the individual DHH and DHHA1 domains is partial, particularly to DHHA1 (the C-terminal domain of DUF4130 shows typical fold decay). None of the catalytic residues in the aforementioned proteins are conserved suggesting that DUF4130 is unlikely to be involved in catalysis. The region of the nucleotide binding site shows some similar features and partial conservation pointing that DUF4130 could be involved in nucleotide/nucleic acid binding. This is further supported by the fact that this domain is sometimes found in combination with the Uracil-DNA glycosylase domain. In the predicted complex DUF4130 occupies the same spatial position as the OB fold domain in MiaB, suggesting DUF4130 could be involved in or regulate the substrate binding of its predicted partner.

**Figure 4.**
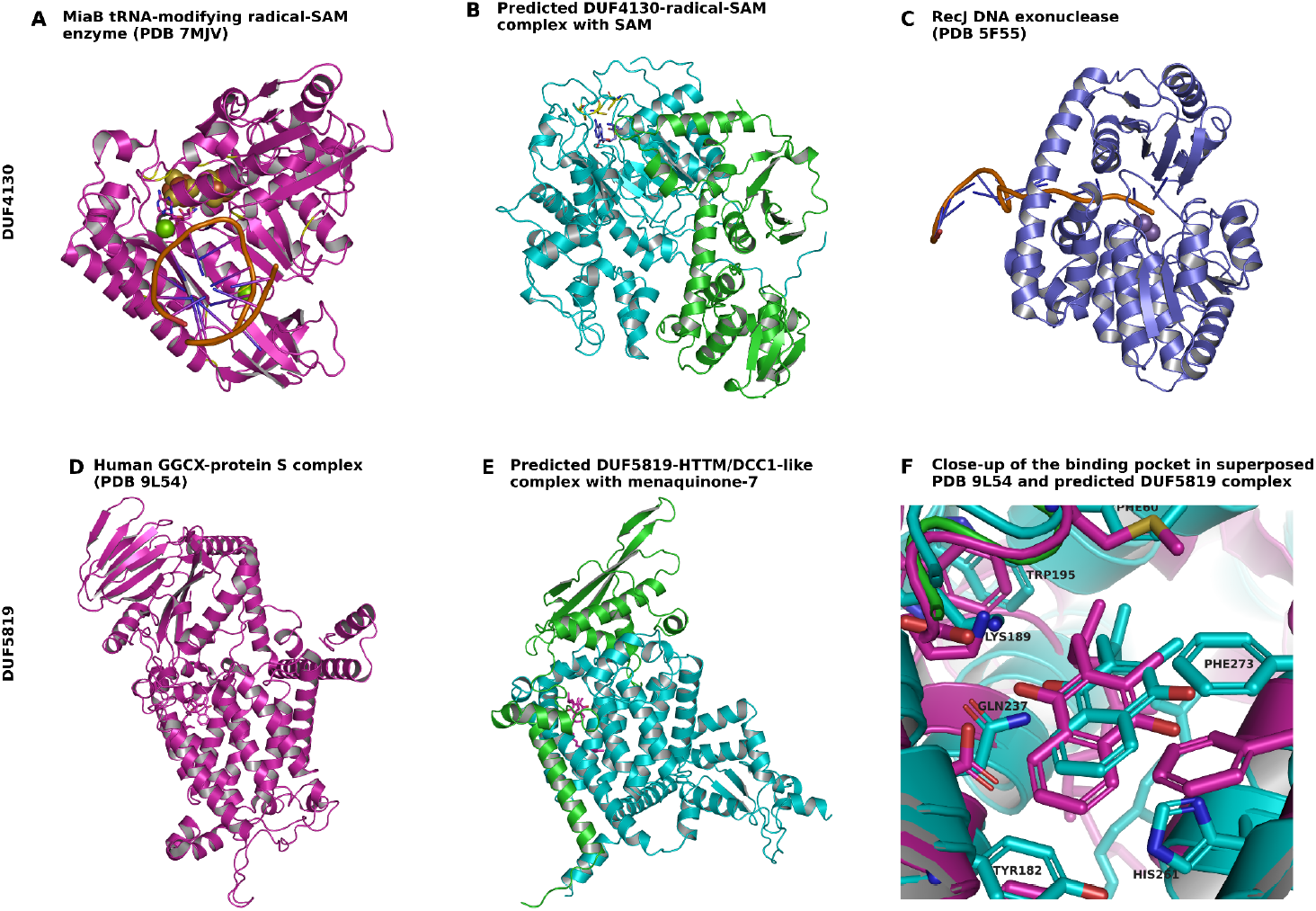
(A-C) Comparison of SAM-bound MiaB, PDB-ID 7MJV: (Esakova *et al*. 2021), the AlphaFold 3-predicted DUF4130 complex with cofolded SAM and DNA-bound RecJ, PDB-ID 5F55: (Cheng *et al*. 2016). Residues 1-42 and 429–704 of RecJ are omitted for visual clarity. Proteins are shown as cartoon, ions as spheres and S-Adenosyl methionine (blue) and the cysteine motif (yellow) as sticks. In (B) the DUF-containing protein is shown in green, while the partner protein is shown in cyan. (D-F) Comparison of vitamin K-dependent gamma-glutamyl carboxylase (PDB-ID 9L54), the AlphaFold 3-predicted DUF5819 complex with cofolded menaquinone-7 hydroquinone (bacterial vitamin K2) and a close-up of the predicted vitamin K-binding pocket. Proteins are shown as cartoon and in (E), the DUF-containing protein is shown in green, while the partner protein is shown in cyan. (D) shows the substrate peptide in yellow as cartoon. (F) is a superposition of PDB-ID 9L54 and the predicted cofolded complex, with important active side residues shown as sticks.

The second example is a DUF5819 protein SAMN04488112_102126 (UniProtKB: A0A1G6IAS5) and SAMN04488112_102125 (UniProtKB: A0A1G6IBF2) from Melg*hirimyces thermohalophilus*. These genes are also adjacent. Their STRING association had a combined score of 959. The predicted complex reached strict confidence, with an interface ipSAE of 0.806 and an average pLDDT of 85.2. Its total interface area was 3,518 Å^2^, including 2,706 Å^2^ contributed by DUF5819. The DUF5819-containing protein contains an N-terminal transmembrane helix followed by DUF5819 at residues 35-192. Its partner is a 447-residue membrane protein containing an HTTM domain at residues 52-289 and a DCC1-like domain at residues 323-431. The current UniProt name, “predicted thiol-disulfide oxidoreductase YuxK, DCC family” is from TrEMBL and unreviewed. This HTTM domain is also present in the vitamin K-dependent gamma-glutamyl carboxylase (GGCX), an enzyme that carboxylates vitamin-K-dependent blood-clotting and anticoagulant proteins in animals (Schultz 2004, Wu *et al*. 2025). Structure comparison of the predicted complex and the human GGCX in a vitamin K-dependent protein S complex (PDB-ID 9L54 (Wu *et al*. 2025)) showed considerable similarity of the modeled complex to the whole GGCX structure. The transmembrane helix of DUF5819-containing protein aligned well with one of the transmembrane helices of GGCX, whereas the DUF5819 domain occupied the spatial position of the GGCX clamp domains. To examine this further, the complex was predicted again with cofolded menaquinone-7 hydroquinone (bacterial vitamin K2), which showed a placement similar to the human GGCX complex, close to the conserved residues Lys189, Trp195 and Phe273. In human GGCX, substrate recognition involves a clamp-like region, where substrate-propeptide binding induces clamp closure that is proposed to stabilize the enzyme-substrate complex during carboxylation (Wu *et al*. 2025). The bacterial DUF5819 complex overlapped with one side of this clamp-like region and residues Leu59 and Phe60 contacted the predicted vitamin K pocket, but the corresponding second side of the clamp was not present and the substrate channel appeared blocked, while a different opening to the active site was accessible. The bacterial partner also contained additional helices at the C-terminus, which were not present in human GGCX suggesting the predicted complex is likely functionally distinct. The predicted complex also resembled the bacterial vitamin K-dependent carboxylase MloH, in which the HTTM region and a region structurally corresponding to DUF5819 are fused. MloH α-carboxylates an aspartyl residue within a hybrid polyketide-nonribosomal peptide intermediate during malonomycin biosynthesis (Law *et al*. 2018).

## 4 Discussion

We found that combining STRING associations with AlphaFold 3 predictions and domain-level interface analysis can refine DUF annotations. The L2 model made the all-organism screen feasible by reducing the number of candidate pairs selected for structure prediction. At the 0.5 model-score cutoff, the final screen contained 2,076 DUF families, including 1,027 with at least one liberal-confidence complex and 681 with at least one strict-confidence complex. The 649 liberal-confidence and 348 strict-confidence complexes with functionally annotated non-DUF partners provide informative candidates for inferring possible DUF functions from their predicted interaction partners. The retrospective comparison showed that predicted partner context was consistent with later Pfam annotations for some families, although the degree of agreement varied. By comparison, although AFDB contained 212,084 DUF-containing heterodimer predictions, only 321 DUF families had a liberal-confidence member and 80 had a strict-confidence member, indicating substantially lower family-level diversity. However, the L2 model should not be considered a general PPI classifier. Its features are correlated channels of STRING and the labels represent AlphaFold 3 prediction outcomes rather than experimentally validated interactions. Adding AFDB heterodimer data did not improve the performance of the L2 model, likely because of the differences in dataset composition and structure-prediction methods. XGBoost was evaluated as an alternative classifier, but not used as final model because the small performance improvement did not justify the reduced interpretability (**Table S2**) (Chen and Guestrin 2016). In general, predicting the output of another computational model creates an inherent performance ceiling and limits generalizability without additional independent biological information. Variation between AlphaFold 3 prediction runs was not assessed because only one diffusion sample and a single seed were used to limit computational cost and enable reproducibility.

Taxonomic recurrence was not used for the logistic regression model because it depends too much on specific species, clustering and STRING thresholds, limiting portability. Also in the final results, liberal-confidence fractions were similar across most recurrence bins, indicating that recurrence alone was not a strong predictor once candidates had passed the L2 prescreening. An increase was only observed among the most recurrent modules, suggesting that only very broadly conserved associations may provide additional support. Genomic proximity is partly represented in STRING through genomic-context evidence such as gene neighborhood, gene fusion and phyletic co-occurrence, but locus distance still separated high- and low-yield prediction groups, suggesting that local gene organization added useful context beyond the final STRING-based candidate selection. In addition, gene distance was estimated from locus-tag numbering and should therefore be regarded only as an approximate measure of genomic proximity. The low confidence yield of the fusion-supported cohort indicates that fusion evidence alone was not sufficient to identify confidently predicted heterodimers. Also several complexes from the other cohorts, including the DUF5819 example, resembled proteins that occur as fusions in other organisms despite lacking a STRING fusion score.

The DUF4130 interaction suggests that the complex is catalyzing radical-SAM dependent nucleic acid modification, with DUF4130 probably regulating nucleotide/nucleic acid binding specificity. The narrow channel makes single-stranded nucleic acid binding more likely, but the specific catalyzed modification and nucleic acid-type is not possible to determine without further experimental analysis.

The DUF5819 prediction suggests the complex forms a bacterial system for vitamin K-dependent carboxylation, with DUF5819 possibly playing a role in the substrate recognition mechanism. However, the blocked substrate channel to the active site and presence of an alternative opening suggest that the substrate specificity and recognition may differ from human GGCX.

There are still several limitations remaining. AlphaFold predictions do not establish interaction in vivo, protein complex stoichiometry or binding affinity and remain subject to hallucinations like steric clashes and impossible chain paths (Hou *et al*. 2023). The calculated confidence metrics and model predictions are also dependent on protein and interface size (Dunbrack 2025, Genz *et al*. 2025). Small interfaces may be unstable or nonspecific and membrane proteins can form apparently confident interfaces between exposed hydrophobic surfaces. Additionally, ligands, cofactors, additional interaction partners, post-translational modifications and alternative binding orientations may also be absent from the predicted complex.

Network analysis of DUF-containing interactions could provide additional functional information but was limited here because most structurally supported DUF proteins had only one distinct predicted partner. Module-based clustering may also have excluded biologically informative interactions while retaining structurally similar or redundant examples. Future screens could improve candidate diversity by grouping closely related Pfam families and structurally similar proteins before selecting representatives. Very small proteins and proteins lacking informative functional or structural features could also be excluded because their predicted complexes may be more susceptible to overconfident AlphaFold prediction (Orr and Bateman 2026).

Together, the resulting dataset provides a selected set of protein-protein interaction candidates. Among complexes in which the DUF forms part of the interface, strict-confidence predictions represent the strongest candidates for stable interactions, followed by liberal-confidence predictions, although definitive interaction and functional assignments still require experimental validation. The DUF4130 and DUF5819 examples demonstrate how these predictions can narrow possible functions and guide subsequent experimental investigation probing their predicted protein-protein and protein-nucleic acid complexes.

## Acknowledgements

AI-assisted coding tools (Codex, using GPT-5.6 Terra and Sol) were used to assist code development and debugging. The resulting code was reviewed by the authors.

## Data availability

The data underlying this article are available in Zenodo at https://doi.org/10.5281/zenodo.21875362.

The code used for analysis and figure generation is available at https://github.com/linoriep/Proteome-scale-structure-prediction-of-DUF-containing-protein-protein-interactions.

