## Supplementary Data for "Large-scale structure prediction of DUF-containing protein-protein interactions"

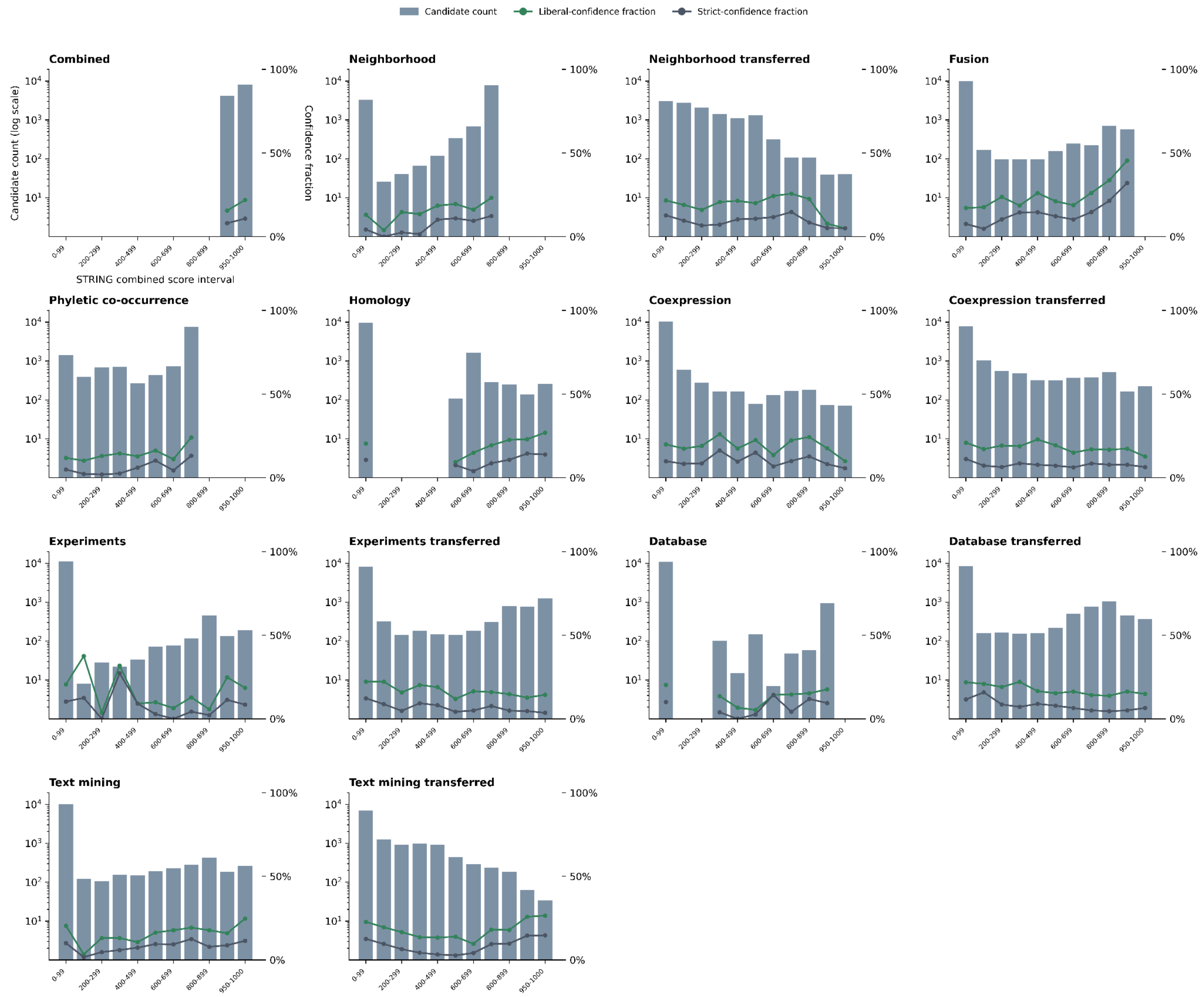


**Figure S1.** Structural-confidence yield across STRING evidence channels in the final L2-selected cohort. Bars show the number of predictions in each score interval. Green and dark grey lines show the fractions meeting the liberal- and strict-confidence criteria, respectively.

**Table S1.** Standardized coefficients of the final L2-regularized logistic regression model.

| Variable | Definition | Standardized Coefficient |
|---|---|---|
| Combined score | Overall STRING association score | 0.366 |
| Neighborhood | Conserved genomic-neighborhood evidence | 0.502 |
| Neighborhood transferred | Neighborhood evidence transferred by orthology | 0.031 |
| Fusion | Gene-fusion evidence | 0.127 |
| Co-occurrence | Phyletic-profile co-occurrence | 0.616 |
| Homology | Sequence-homology score | 0.206 |
| Coexpression | Direct coexpression evidence | 0.058 |
| Coexpression transferred | Coexpression transferred by orthology | -0.083 |
| Experiments | Direct experimental evidence | 0.149 |
| Experiments transferred | Experimental evidence transferred by orthology | -0.046 |
| Database | Curated-database evidence | 0.033 |
| Database transferred | Database evidence transferred by orthology | -0.202 |
| Text mining | Literature co-mention evidence | -0.070 |
| Text mining transferred | Text-mining evidence transferred by orthology | -0.050 |

**Table S2.** Five-fold stratified cross-validation performance of the L2-model with additional AFDB heterodimer training data, an XGBoost model and baseline comparisons based on the combined STRING score, neighbourhood STRING channel and co-occurrence STRING channel. Model performance was evaluated using five-fold stratified cross-validation. XGBoost used 100 trees, a maximum depth of 2, a learning rate of 0.05, a minimum child weight of 12, row subsampling of 0.85 and feature subsampling of 0.85.

| Training Configuration | AP | AUROC |
|---|---|---|
| L2 Model STRING900/taxa10 (default) | 0.3716 | 0.8003 |
| L2 Model STRING900/taxa10 + AFDB heterodimer data | 0.3406 | 0.7871 |
| XGBoost STRING900/taxa10 | 0.4002 | 0.8042 |
| Combined STRING score | 0.1419 | 0.5245 |
| Neighbourhood STRING channel | 0.2440 | 0.7373 |
| Co-occurrence STRING channel | 0.3163 | 0.7806 |

**Table S3.** Ordered keyword rules used to classify partner-protein functions.

| Priority | Partner-function category | Keyword rule |
|---|---|---|
| 1 | translation / RNA | ribosom, translation, tRNA, rRNA, mRNA, ribonuclease, RNA polymerase, elongation factor, initiation factor, termination factor, DEAD-box, ribosome-binding |
| 2 | transport / membrane | transporter, transport, permease, ABC transporter, ABC-type, symporter, antiporter, channel, porin, pump, efflux, membrane, lipoprotein, secretion, export, import, uptake, translocase, periplasm, envelope, outer membrane, inner membrane |
| 3 | cell envelope / division | cell wall, peptidoglycan, murein, lipopolysaccharide, capsule, division, septum, Fts proteins, sporulation, spore, flagellum, pilus, fimbriae, biofilm, lysis, rod shape, MreB, RodZ |
| 4 | DNA / chromosome | DNA, DNA replication, DNA repair, recombination, helicase, topoisomerase, polymerase, nuclease, chromosome, gyrase, mismatch repair, RecA, Uvr proteins, DNA-binding, SOS response, primase, plasmid, integrase, transposase |

| | | |
|---|---|---|
| 5 | metabolism / enzyme | dehydrogenase, synthetase, synthase, kinase, phosphatase, transferase, hydrolase, oxidase, reductase, isomerase, lyase, ligase, carboxylase, aminotransferase, biosynthesis, catabolic, decarboxylase, mutase, peptidase, aldolase, enzyme, ATP binding, racemase, esterase, lipase, deaminase, redox, thioredoxin, flavin, heme |
| 6 | regulation / signaling | regulator, transcriptional, response regulator, sensor, two-component, sigma, anti-sigma, repressor, activator, cyclic, signal, modulates, competence, HTH, helix-turn-helix |
| 7 | stress / defense | stress, toxin, antitoxin, resistance, detox, oxidative, heat shock, cold shock, chaperone, protease, restriction-modification, CRISPR, defense, virulence, antibiotic, phage, prophage, bacteriophage, capsid, terminase, baseplate, tail, sheath, portal, holin, immunity |
| 8 | unknown-function protein | DUF, unknown, uncharacterized, uncharacterised, hypothetical, putative protein, conserved protein, protein of unknown function |
| 9 | other / unclear | no category or unknown-function rule matched |

**Table S4.** Retrospective comparison of DUF families and partner function relation, in the L2-screen (Pfam 38.0) with Pfam 38.2 annotations.

| Pfam ID | Former DUF | Pfam 38.2 name | Agreement | Partner-context evidence |
|---|---|---|---|---|
| PF01345 | DUF11 | CLIPPER | Broader process | Type IX secretion and cell-envelope partners are consistent with an envelope-associated context but do not define CLIPPER function. |
| PF01732 | Mycop_pep_DUF31 | Myco_protease | Specific | Immunoglobulin-binding and immunoglobulin-blocking partners are consistent with a Mycoplasma immunoglobulin protease system. |
| PF02594 | DUF167 | PF1765-like | No clear relationship | Partners do not indicate a specific relationship to the PF1765-like family assignment. |
| PF04685 | DUF608 | GH116_catalytic | Specific | Beta-glucosidase and GH116-containing partners are consistent with the GH116 catalytic-region assignment. |
| PF06075 | DUF936 | CORD_N | No clear relationship | The single partner does not indicate a relationship to the CORD N-terminal assignment. |
| PF06672 | DUF1175 | MagC | Broader process | Alpha-2-macroglobulin and YfaP-associated partners are consistent with an envelope or protease-defense system but not the specific MagC function. |
| PF07849 | DUF1641 | HMP | Broader process | Formate dehydrogenase and oxidoreductase partners are consistent with a membrane respiratory-complex context without defining the HMP role. |
| PF09836 | DUF2063 | HvfC_N | Specific | BufA1- and MNIO-associated partners are consistent with the HvfC/BufC role in bufferin precursor modification. |
| PF09906 | DUF2135 | YfaP_C | Broader process | Alpha-2-macroglobulin and MagC partners are consistent with the same envelope or protease-defense system without defining YfaP function. |
| PF09979 | DUF2213 | Pa193_gp24 | Broader process | Phage structural and packaging partners indicate a phage-assembly context but do not specifically identify the gp24 scaffolding function. |
| PF10022 | DUF2264 | BcGHD | No clear relationship | Predominantly self-annotations and ABC-transporter partners do not indicate beta-D-glucuronate dehydratase function. |
| PF10048 | DUF2282 | BufA1 | Specific | HvfC/BufC-associated partners are consistent with BufA1 as a bufferin metallophore precursor. |

| | | | | |
|---|---|---|---|---|
| PF10240 | DUF2464 | MABP | Specific | VPS37B, TSG101 and VPS28 partners place the domain in the ESCRT-I context associated with MABP/MVB12. |
| PF10709 | DUF2511 | YebY | Specific | The YebZ partner places YebY in the named AZY copper-uptake system. |
| PF11175 | DUF2961 | GH172_2nd | Broader process | Glycosyl hydrolase and carbohydrate-binding partners are consistent with carbohydrate processing without identifying GH172 specifically. |
| PF11816 | DUF3337 | Ubl_WDR48-Bun107 | Specific | USP1 and Fanconi-pathway partners are consistent with the WDR48/Bun107 ubiquitin-associated context. |
| PF11863 | DUF3383 | E217_gp31_N | Broader process | Dit-like and other phage-tail partners indicate a tail-assembly context but do not specifically identify the gp31 N-terminal region. |
| PF11958 | DUF3472 | BPSS1860 | Broader process | An outer-membrane starch-binding partner is consistent with a surface-associated context without identifying BPSS1860 function. |
| PF11984 | DUF3485 | EpsI | Specific | Exosortase and TPR-rich partners are consistent with an extracellular-polysaccharide system associated with methanolan biosynthesis EpsI. |
| PF12532 | DUF3732 | LmuB_C | No clear relationship | Diverse ABC and defense-associated partners do not indicate a specific relationship to LmuB. |
| PF12996 | DUF3880 | YkvP_N | Broader process | Cell-wall glycosyltransferase partners are consistent with an envelope-associated process without specifically identifying YkvP. |
| PF13835 | DUF4194 | JetB | Broader process | DNA-repair and defense-associated partners indicate an antiphage-defense context but do not specifically identify JetB. |
| PF14092 | DUF4270 | BamH | No clear relationship | Galactose-oxidase and Kelch-repeat partners suggest a beta-propeller context, but no clear relationship to BamH was identified. |
| PF14298 | DUF4374 | XusB | Specific | TonB-dependent iron-uptake receptors are consistent with the xenosiderophore-binding XusB assignment. |
| PF14332 | DUF4388 | PatA_N | Broader process | GGDEF and response-regulator partners are consistent with a bacterial signaling context but do not specifically identify the PatA N-terminal domain. |
| PF16090 | DUF4819 | Cic | Specific | The Ataxin-1 partner is consistent with the known protein context of the capicua homolog Cic. |
| PF16324 | DUF4960 | SGBP_levan-binding | Specific | SusC(lev), SusD and beta-fructofuranosidase partners are consistent with a levan-utilization system. |
| PF16332 | DUF4962 | DSE-like_N | Broader process | A heparinase partner is consistent with polysaccharide processing without establishing dermatan-sulfate epimerase activity. |
| PF16472 | DUF5050 | LRP1-like_beta_prop | Broader process | EGF-like and extracellular partners are consistent with an LRP-like architecture without identifying the beta-propeller’s specific function. |
| PF17270 | DUF5336 | MAP_0900 | No clear relationship | No informative partner annotation was available for comparison with the 34 kDa antigenic-protein assignment. |
| PF17384 | DUF150_C | RimP_C | Specific | Ribosomal-protein partners are consistent with the ribosome-assembly context of RimP. |
| PF17825 | DUF5587 | SHOC1_ATPase | No clear relationship | The only partner repeats the SHOC1 assignment and provides no independent functional context. |
| PF19528 | DUF6056 | GT_STM0557_N | Specific | A phage glycosyltransferase partner is consistent with the glycosyltransferase-domain assignment. |

| | | | | |
|---|---|---|---|---|
| PF19557 | DUF6079_1st | BrxC-PglY_N | Broader process | DNA methylase and ATPase partners indicate a defense-system context but do not distinguish BrxC from PglY. |
| PF24346 | DUF7507 | CLIPPER_2 | Broader process | Type IX secretion and CLIPPER-containing partners indicate the expected cell-surface context but do not specifically establish CLIPPER 2 function. |
| PF25372 | DUF7885 | LRR_FBXL15 | Specific | SKP1 and cyclin-associated partners are consistent with an F-box/LRR protein context. |
| PF26062 | DUF8022 | PccX | Specific | A propionyl-CoA carboxylase carboxyltransferase partner is consistent with the PccX assignment. |
| PF26607 | DUF8189 | Beta-prop_PLL | No clear relationship | Varied phage and defense-associated partners do not establish a relationship to the PLL-like beta-propeller assignment. |
